# The transcriptome of *Schistosoma mansoni* developing eggs reveals key mediators in pathogenesis and life cycle propagation

**DOI:** 10.1101/2021.05.20.444946

**Authors:** Zhigang Lu, Geetha Sankaranarayanan, Kate Rawlinson, Victoria Offord, Paul J. Brindley, Matt Berriman, Gabriel Rinaldi

## Abstract

Schistosomiasis, the most important helminthic disease of humanity, is caused by infection with parasitic flatworms of the genus *Schistosoma*. The disease is driven by the eggs laid by the parasites and becoming trapped in host tissues, followed by inflammation and granuloma formation. Despite abundant transcriptome data for most developmental stages of the three main human-infective schistosome species, i.e. *Schistosoma mansoni, S. japonicum* and *S. haematobium*, the transcriptomic profiles of developing eggs remain largely unexplored. In this study, we performed RNAseq of *S. mansoni* eggs laid *in vitro* during early and late embryogenesis (days 1-3 and 3-6 post-oviposition, respectively). Analysis of the transcriptomes identified hundreds of up-regulated genes during the later stage, including venom allergen-like (VAL) proteins, well-established host immunomodulators, and genes involved in organogenesis of the miracidium larva. In addition, the transcriptomes of the *in vitro* laid eggs were compared with existing publicly available RNA-seq dataset from *S. mansoni* eggs collected from the livers of murine hosts. Analysis of enriched GO terms and pathway annotations revealed cell division and protein synthesis processes associated with early embryogenesis, whereas cellular metabolic processes, microtubule-based movement, and microtubule cytoskeleton organization were found enriched in the later developmental time point. This is the first transcriptomic analysis of *S. mansoni* embryonic development, and will facilitate our understanding of infection pathogenesis, miracidia development and life cycle progression of schistosomes.

## Introduction

Schistosomes are parasitic flatworms that infect more than 250 million people worldwide, mainly in Low and Middle-Income Countries (Toor et al., 2018). There is only a single effective drug (praziquantel), and an ongoing threat of drug resistance emerging (Crellen et al., 2016). While adult worms can dwell within the blood vessels of a human host for years, it is their eggs rather than the worms themselves that drive pathology (Pearce and MacDonald, 2002). It is estimated that one pair of *Schistosoma mansoni* adult worms can lay >300 eggs per day (Cheever et al., 1994). Once the eggs are laid by the worms in the mammalian bloodstream, about half migrate, over the course of 6 days, through the endothelium of blood vessels, across the epithelium of the gut and are released into the intestinal lumen (Jourdane and Théron, 1987; Schwartz & Fallon, 2018; Costain et al., 2018). Within the lumen, the eggs take ∼4 hours to transit along the intestinal tract inside faecal material (Wang et al.,1999). The egg excreted into water hatches a miracidium larva that must infect a freshwater snail to continue the life cycle. The remaining 50% of eggs are swept around the body by the bloodstream and become trapped in the host’s liver, intestines and spleen, where they induce immune responses, severe inflammation and granuloma formation (Ross et al., 2002). The granuloma surrounding the trapped egg consists of an immune cellular complex that includes macrophages, lymphocytes and eosinophils (Pearce and MacDonald, 2002). In addition, fibroblasts in the granuloma produce collagen that leads to periportal fibrosis, induction of collateral circulation including varices which, in turn, increase the risk of life-threatening digestive haemorrhage (Gryseels et al., 2006).

The study of the schistosome egg and its interaction with the host is critical to understand not only the pathogenesis associated with the infection, but also parasite strategies to exit the mammalian host to continue the life cycle (Costain et al., 2018; Schwartz and Fallon, 2018). Thus, a myriad of reports have focused on soluble egg antigens (SEA) and several excreted-secreted products, including proteins and glycans that interact with the host tissues inducing immune responses and facilitate the egress of the egg to the external environment (Asahi and Stadecker, 2003; Schramm et al., 2003, 2009; Fitzsimmons et al., 2005; Meevissen et al., 2012). Notably, the immune modulatory roles of egg-specific antigens, such as Omega, Kappa and IPSE have been validated by functional approaches such as shRNA-mediated knock-down (Hagen et al., 2014) and CRISPR-Cas-based genome editing (Ittiprasert et al., 2019). More recently, the molecular and cellular mechanisms involved in granuloma formation have been dissected using a zebrafish model of macrophage dependent granuloma induction (Takaki et al., 2021a). This novel infection model demonstrated that host and parasite molecules play key roles in shaping the granulomatous response and that the response is dependent on the level of egg maturity (Takaki et al., 2021a; Takaki et al., 2021b).

Notwithstanding this progress, few studies in recent years have focused on the development of the miracidia inside the egg capsule (Michaels and Prata, 1968). Jurberg et al. (2009) provided a detailed morphological description of embryogenesis of the miracidia inside the egg capsule while migrating through the host tissue. In addition, these investigators suggested a new staging system for miracidia development that comprises eight discrete stages (Jurberg et al., 2009). No transcriptome analyses underlying this developmental progression have yet been performed, with only a single RNA-seq report of *S. mansoni* eggs isolated from the liver of experimentally-infected hamsters (Anderson et al., 2015). Aiming to address this lack of information, we performed comparative transcriptomics (RNA-seq) on *in vitro* laid eggs of *Schistosoma mansoni* at two time points; early and late embryogenesis. More than 1,300 genes were differentially expressed, including up-regulation in the late development stage of genes associated with organogenesis of the miracidium larva. The investigation revealed transcriptional signatures in developing *in vitro* laid eggs that will facilitate our understanding of the infection pathogenesis, miracidia development and life cycle progression of schistosomes.

## Material and methods

### Ethics statement

The complete life cycle of *Schistosoma mansoni* NMRI (Puerto Rican) strain is maintained at the Wellcome Sanger Institute (WSI) by breeding and infecting susceptible *Biomphalaria glabrata* snails and mice. The mouse experimental infections and other regulated procedures were conducted under the Home Office Project Licence No. P77E8A062 held by GR. All protocols were revised and approved by the Animal Welfare and Ethical Review Body (AWERB) of the WSI. The AWERB is constituted as required by the UK Animals (Scientific Procedures) Act 1986 Amendment Regulations 2012.

### *In vitro* laid eggs

Eggs of *Schistosoma mansoni* laid *in vitro* by cultured adult worms (*in vitro* laid eggs or IVLE) were collected as described (Rinaldi et al., 2012). Briefly, mixed-sex adult worms were recovered from mice by portal perfusion 6 weeks after infection, washed in 1X PBS supplemented with 200 U/ml penicillin, 200 μg/ml streptomycin and 500 ng/ml amphotericin B (ThermoFisher Scientific), transferred to 6-well plates, and maintained in culture in complete Basch media at 37°C in 5% CO2 (Mann et al., 2010). All media components were purchased from ThermoFisher Scientific. The eggs laid by the worms in culture during the first 72 h post-perfusion were recovered. Fifty percent of the eggs were collected at this time for the early embryogenesis samples (D3 eggs), concentrated by gravity, resuspended in 750 µl Trizol reagent, snap frozen and stored at -80°C. The rest of the IVLE were cultured for a further 3 days to allow further development before collection for the late embryogenesis samples (D6 eggs) (Mann et al., 2011). A total number of eggs ranging from 500 to 1000 IVLE were collected at each time point. We performed a separate collection of IVLE, as above, for each of three independent perfusions of adult schistosomes from experimentally infected mice.

### RNA extraction from *in vitro* laid eggs

The frozen IVLE in Trizol reagent were subjected to 3 rounds of freeze thaw cycles by manually transferring the tubes from the water bath at 95°C to dry ice. This procedure enhanced the total RNA yield from the samples. Thereafter, the eggs were transferred to Magnalyser tubes containing ceramic beads, homogenized in FastPrep (FastPrep-24, MP Biomedicals) at setting 6 with two 20-second pulses and incubated for 5 minutes at room temperature. One hundred and fifty µl of chloroform was added to each sample, shaken vigorously for 10 seconds, incubated for 3 minutes at room temperature and centrifuged at 15,000g for 15 minutes at room temperature. The aqueous phase was carefully removed to a clean centrifuge tube, and RNA precipitated by adding an equal volume of 100% ethanol and incubating at -80°C overnight. The samples were centrifuged at maximum speed for 30 minutes at 4°C, the RNA pellet washed in 70% ethanol, air dried and resuspended in nuclease free water. The RNA quality was assessed and quantified using the Bioanalyzer (2100 Bioanalyzer Instrument, Agilent Technologies). Although the overall yield of RNA was modest in quantity, <30 ng total, a high-quality RNA was recovered from each replicate sample **(Suppl. Figure S1A**).

### Library preparation and sequencing

Given the minimal amount of RNA obtained from IVLE, the single cell Smart-Seq2 protocol was adapted for low-input RNA library preparation (Picelli et al., 2014). Two different amounts of input RNA, 12 ng and 3 ng, for each sample and its replicates were prepared. Poly A-tailed mRNA was enriched using Dynabead Oligo(dT)20 at 5mg/ml concentration 5 mg/mL in 1X PBS (pH 7.4) from mRNA DIRECT kit (61011). Beads were washed using wash buffer with 10mM Tris-HCl pH 7.5, 150 mM LiCl, 1mM EDTA pH 8.0, 0.1% w/v LiDS in nuclease free water and RNA eluted from beads using the 10 mM Tris-HCl pH 8.0 at 75°C for 2 minutes. The poly A-enriched RNA was reverse transcribed by SmarSeq2 method as described (Picelli et al., 2014) with 10 RT cycles and cDNA further amplified using ISPCR primers with 10 or 11 rounds of PCR cycles. Dual indexed sequencing libraries were made out of 5 ng cDNA from the above preparations using Illumina Nextera library preparation kit according to manufacturer’s instructions (Illumina). Quality checked and equimolar pooled libraries were sequenced in a HiSeq 4000 Illumina system. Sequence data were deposited with the study number ERP128933, and accession numbers for each sample is shown in **Supp. Table S1**.

### Mapping of RNA-seq reads and gene counting

Sequence reads from eggs isolated from the liver of experimentally-infected hamsters 6 to 8 weeks after infection (‘liver eggs’) were obtained from published data (Anderson et al., 2015). The reads were mapped to *S. mansoni* v7 genome (WormBase Parasite WBPS14) using Hisat 2.1.0 (Kim et al., 2015) due to unequal read lengths generated on Roche 454. Sequence reads for IVLE were mapped using STAR 2.5.0a (Dobin et al., 2013) with the option --alignIntronMin 10. Counts per gene were summarised with FeatureCounts v1.4.5-p1 (Liao et al., 2014) based on the exon feature, using the annotation from WormBase Parasite WBPS14 (https://parasite.wormbase.org/).

### Differential gene expression analysis

Raw read counts from both liver eggs (Anderson et al., 2015) and IVLE samples (this study) were combined and used as input for DESeq2 v1.26.0 (Love et al., 2014). The Pearsons’s correlations between replicates were examined. For differential expression analysis, we set cooksCutoff=TRUE to remove extreme outlier genes. Log-transformed count data were used for principle component analysis (PCA) and calculating the euclidean distance between samples.

### Gene Ontology enrichment analysis of differentially expressed genes

Gene Ontology (GO) annotation for *S. mansoni* genes were obtained by running InterProScan v5.25 (Mitchell et al., 2019). Enrichment analysis of differentially expressed genes (DEGs) were performed using topGO v2.38.1 (Alexa and Rahnenfuhrer, 2020), with 5 nodes and the weight 01 method. GO terms with FDR < 0.05 were considered as significantly enriched. Gene product descriptions were obtained using the Biomart function on WormBase Parasite (https://parasite.wormbase.org/).

### Enrichment of Pfam family and InterPro domains

The annotations of Pfam family and InterPro domain in *S. mansoni* gene products were obtained from InterProScan v5.25. Analysis of functional enrichment in DEGs were conducted via Fisher’s Exact test followed by a P-value correction using the Benjamini-Hochberg procedure. Terms with FDR < 0.05 were considered as significant.

### KEGG pathway mapping

Mapping of *S. mansoni* gene products to the KEGG pathway database was performed on the KAAS server (https://www.genome.jp/kegg/kaas/) using the GHOSTX program and BBH method. The significance of DEGs enrichment in pathways was assessed using Fisher’s Exact test and resulting P-values were adjusted using the Benjamini-Hochberg procedure for each of KEGG categories 1-5 (https://www.genome.jp/kegg/pathway.html; excluding pathways for prokaryotes, yeast, and plant). Pathways with FDR < 0.05 were considered as significant. The scripts used for functional enrichment (GO, Pfam, InterPro, KEGG) analysis can be accessed at https://github.com/zglu/Gene-function-enrichment.

## Results

### Transcriptional signatures underlie the *in vitro* development of *Schistosoma mansoni* eggs

Aiming at studying the transcriptional profiles associated with the *in vitro* development of eggs, we performed RNA-seq from early embryogenesis samples (1–3 days post oviposition (D3) and late embryogenesis samples (4–6 days oviposition (D6)) **(Figure 1A)**. Most D3 eggs belonged to stages I (non-visible embryo under the light microscope) and II (visible embryo as a clear central disk that occupies one third of the egg) (Michaels and Prata, 1968; Jurberg et al., 2009) **(Figure 1B** and **Suppl. Figure S2A)**. By six days after worm collection (D6 eggs), the eggs were further developed and had increased in size by one third, as previously reported (Jurberg et al., 2009). In addition, 40–45% of the D6 eggs had progressed to stages III (enlarged embryo that occupies two thirds of the egg length), IV (embryo occupying almost the entire egg), or V (fully mature miracidium inside the eggshell before hatching, some motile miracidia) (Michaels and Prata, 1968; Jurberg et al., 2009) **(Figure 1C** and **Suppl. Figure S2B**; **Suppl. Video S1)**.

**Figure 1.**
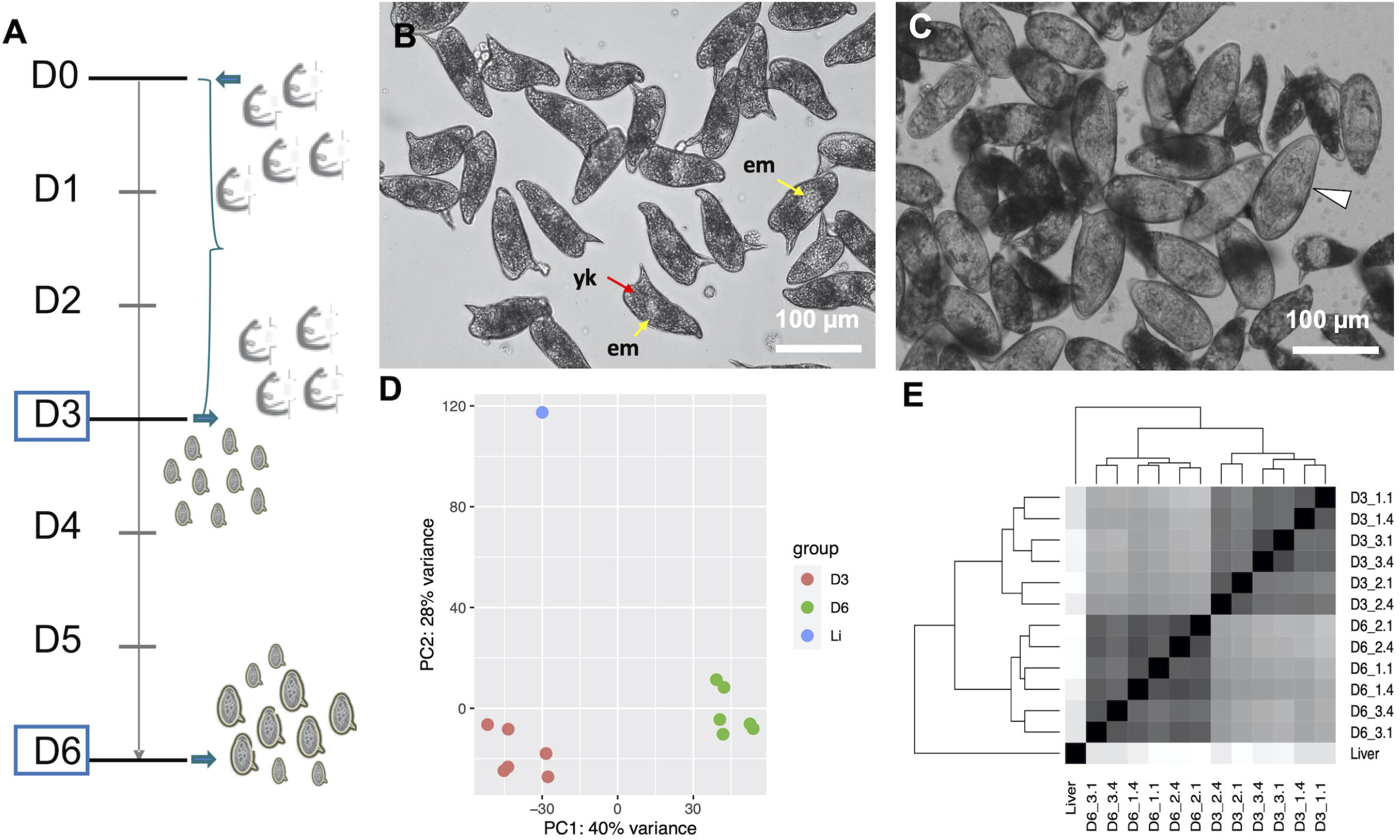
**A**. Timeline depicting the experimental design. On day 0 (D0) adult worms perfused from experimentally infected mice were washed and placed in culture for three days. The worms laid eggs, i.e. *in vitro* laid eggs (IVLE), for three days (bracket). On day 3 (D3) the worms were removed from the culture, and half of the IVLE were collected for RNAseq - D3, early embryogenesis sample. The rest of the IVLE were kept in culture for three more days, and on day 6 (D6) they were collected for RNAseq - D6, late embryogenesis sample. **A, B**. Representative micrographs of 3 days old- (A) and 6 days old- (B) *in vitro* laid eggs (IVLE). Scale bar: 100 μm. *em*, embryo (yellow arrow); *yk*, yolk (red arrow); white arrowhead, fully developed egg containing the mature miracidium. **C**. Clustering of egg samples using Principal Component Analysis. D3: D3 IVLE; D6: D6 IVLE; Li: liver eggs **D**. Clustering of egg samples using Sample Distance Matrix. Samples’ names are described as indicated in Supp. Table S1.

The quality of the total RNA extracted from IVLE was much higher than that of the highly-degraded RNA usually obtained from eggs collected from the liver of experimentally-infected mice **(Suppl. Figure S1B)**. On the other hand, the total RNA yield isolated from ∼500–1000 IVLE was <30 ng. Therefore, we decided to adapt the Smart-Seq2 protocol originally designed for single-cell RNA-seq (Picelli et al., 2014) to produce high-quality RNA-seq libraries from 3 or 12 ng of input RNA. We obtained 0.2–3.3 million RNA-seq raw reads, and 75% of them have >2x reads than the published liver egg sample from which 1000 ng of polyA^+^ RNA was used for RNA-seq (Anderson et al., 2015) (0.36 million reads; **Suppl. Table S1**). All replicates of IVLEs showed good correlations (Pearson’s r > 0.83; **Suppl. Table S2**), and from both Principal Component Analysis (PCA) and Sample Distance Matrix analyses, the samples clustered according to their developmental stage (**Figure 1D, E**).

### *In vitro* developed eggs are transcriptionally distinct from eggs collected from the host

Major transcriptomic differences were evident among the three egg samples; the two IVLE samples—early embryogenesis (D3), late embryogenesis (D6)—and the liver-collected eggs. However, the gene expression profiles of D3 and D6 IVLE partially overlapped suggesting a progression in the transcriptome changes during the egg development **(Figure 2A)**. On the other hand, the transcriptome of liver eggs was distinct to that of IVLE; 240 and 832 genes were significantly up- and down-regulated, respectively, in D3 IVLE compared to liver eggs; 645 and 708 genes were significantly up- and down-regulated, respectively, in D6 IVLE compared to liver eggs (Padj < 0.01 & fold-difference >2; **Figure 2**; **Supp. Figure S3; Suppl. Tables S3** and **S4**). Strikingly, the expression of well-described immunomodulatory egg-specific genes, including *omega-1* (Smp_345790) and *kappa-5* (Smp_335490 & Smp_344300) (Hagen et al., 2014; Ittiprasert et al., 2019) was significantly higher in liver eggs compared to IVLE **(Figures 2B, C** and **3**; **Suppl. Table S4)**. These two egg-specific genes showed more than a 30-fold increase in expression at D6 compared to D3 **(Figure 2)**. In addition, expression of the putative major egg antigen (Smp_302280), extensively studied as a key component in the T-cell-mediated response during the granuloma formation (Stadecker and Hernandez, 1998), significantly increased during the *in vitro* egg development **(Suppl. Table S4)**. However, the overall expression of this gene was the highest in the liver eggs compared to both samples of IVLE **(Suppl. Table S4)**.

**Figure 2.**
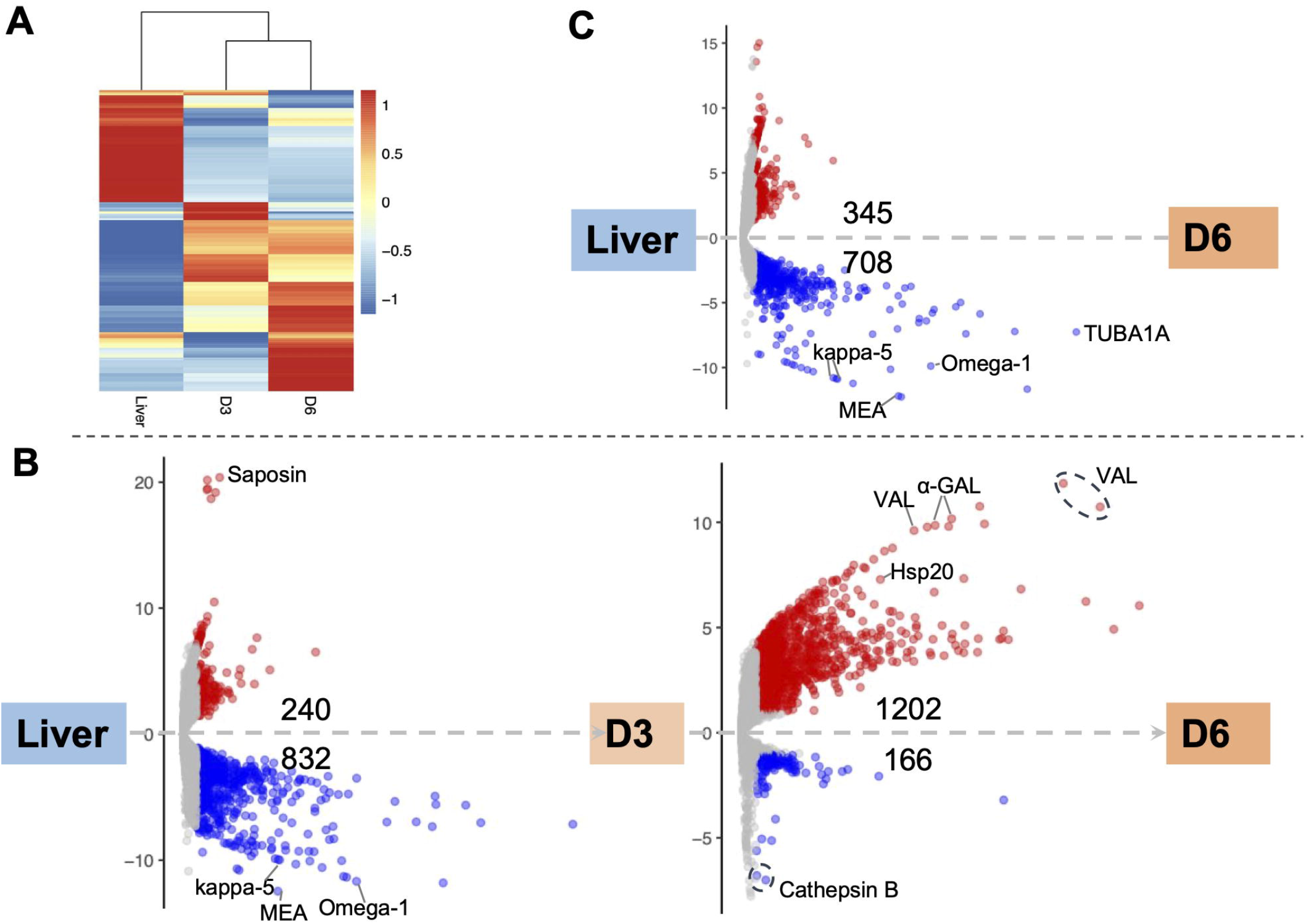
Differential gene expression among egg samples. **A**. Hierarchical clustering showing divergent transcriptomic signatures among the three samples. Liver: liver eggs; D3: D3 IVLE; D6: D6 IVLE. **B**. Volcano plots showing differentially expressed genes (DEGs) in D3 IVLE compared to liver eggs (left) and in D6 IVLE compared to D3 IVLE (right). Highlighted genes: Saposin (Smp_105420), kappa-5 (Smp_335470 & Smp_335480), MEA (major egg antigen - Smp_302280), Omega-1 (Smp_334170), Hsp20 (Smp_302270), VAL (venom-allergen like protein - Smp_070250, Smp_176180 & Smp_120670), α-GAL (Alpha-N-acetylgalactosaminidase - Smp_247750 & Smp_247760), Cathepsin B (Smp_067060 & Smp_103610). **C**. Volcano plot showing DEGs in D6 IVLE compared to liver eggs. Highlighted genes: kappa-5 (Smp_335470 & Smp_335480), MEA (major egg antigen - Smp_302350), Omega-1 (Smp_334170), TUBA1A (Tubulin alpha-1A chain - Smp_090120) In the volcano plots the *x-axes* represent -log10Padj values and the *y-axes* represent log2FoldChange. The volcano plots are available as interactive charts at http://schisto.xyz/IVLE/.

### Miracidium-enriched genes up-regulated in late egg development

Further pairwise analysis between both IVLE transcriptomes, showed 1202 up- and 166 down-regulated genes in D6 compared to D3 eggs **(Figure 2B**; **Suppl. Table S4)**, indicating an overall upregulation of gene expression as embryogenesis proceeded. Out of the 1202 upregulated genes in D6, 524 are among marker genes for different cell types previously defined by single cell RNA-seq in schistosomula, the first intra-mammalian stage (Diaz Soria et al., 2020). Of these, 53.8% (282/524) belong to neurons, 18.7% (98/524) to muscles, 18.9% (99/524) to the parenchyma tissue, and 8.6% (45/524) to germinal cells **(Suppl. Tables S4** and **S5)**.

Genes involved in the interaction between the miracidium and the snail are among the top 10 genes with the highest fold change difference between D6 and D3 eggs **(Suppl. Table S4)**. For example, genes encoding venom allergen-like (VAL) proteins were upregulated in D6 eggs **(Figure 3)**, including VAL 9 (Smp_176180), VAL 5 (Smp_120670), and VAL 15 (Smp_070250) **(Figure 2B**; **Suppl. Table S4)**. Functional analysis based on Gene Ontology (GO) and KEGG Pathways revealed biological processes and molecular functions associated with differentially-expressed genes among the three egg samples **(Figure 4**; **Suppl. Table S6)**. The transcriptome of D3 compared to D6 eggs or liver eggs, showed an enrichment for DNA replication, cell cycle, ribosome biogenesis and RNA translation **(Figure 4)**. This is consistent with the cell division and protein synthesis which are critical processes in the early developing embryo (Jurberg et al., 2009). In contrast, up-regulated genes in D6 eggs were associated with processes associated with movement and signalling, e.g. microtubule-based movement, signal transduction, and GPCR signalling pathways (FDR<0.05; **Figure 4**; **Suppl. Table S6**). The microtubule motor activity associated with the later developmental time point may be related to the development of the ciliary plates of the miracidium that enable swimming.

**Figure 3.**
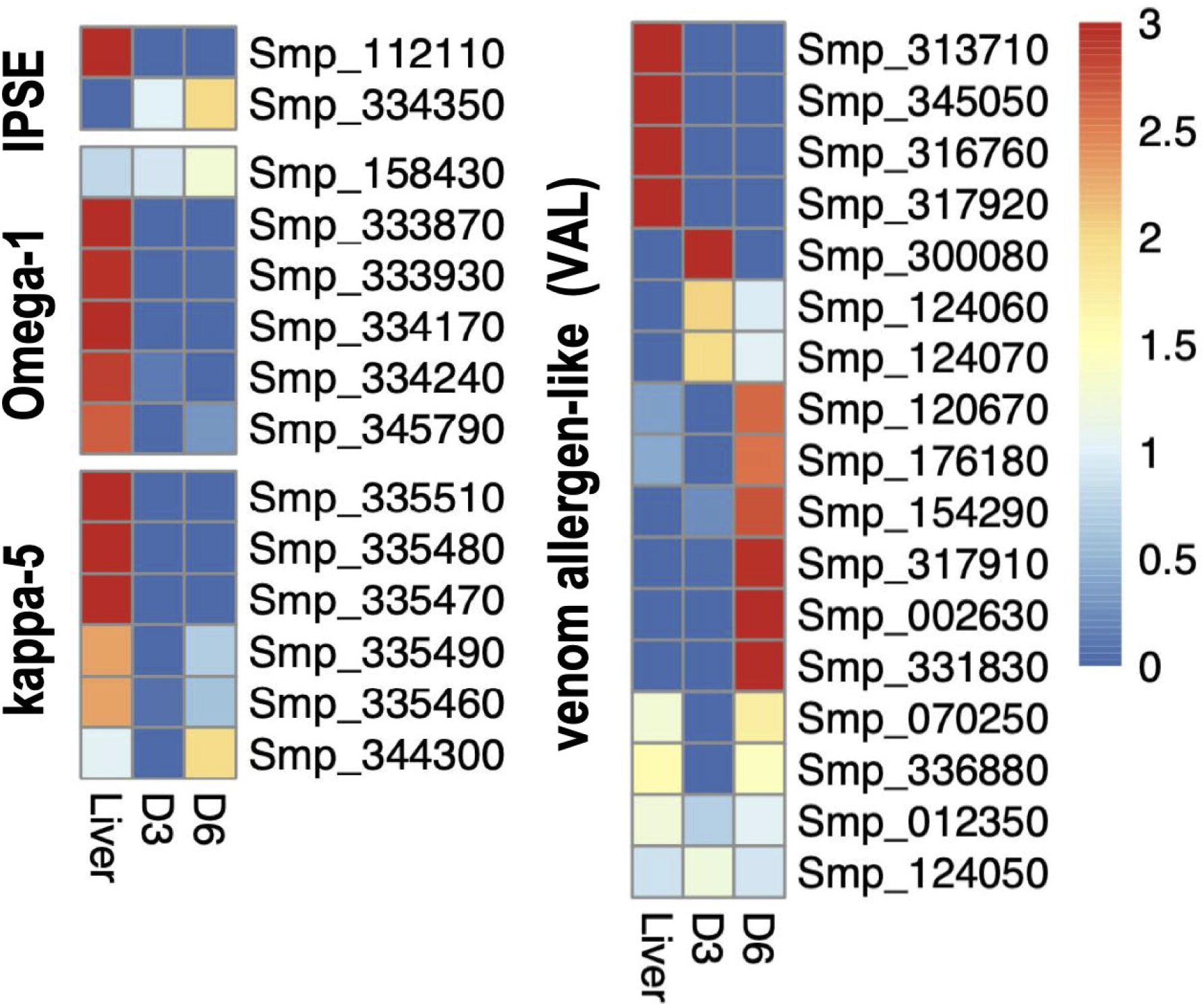
Heatmaps showing the relative gene expression of the egg antigens IPSE, Omega-1, kappa-5 and VALs in liver, D3 and D6 IVLE, as indicated.

**Figure 4.**
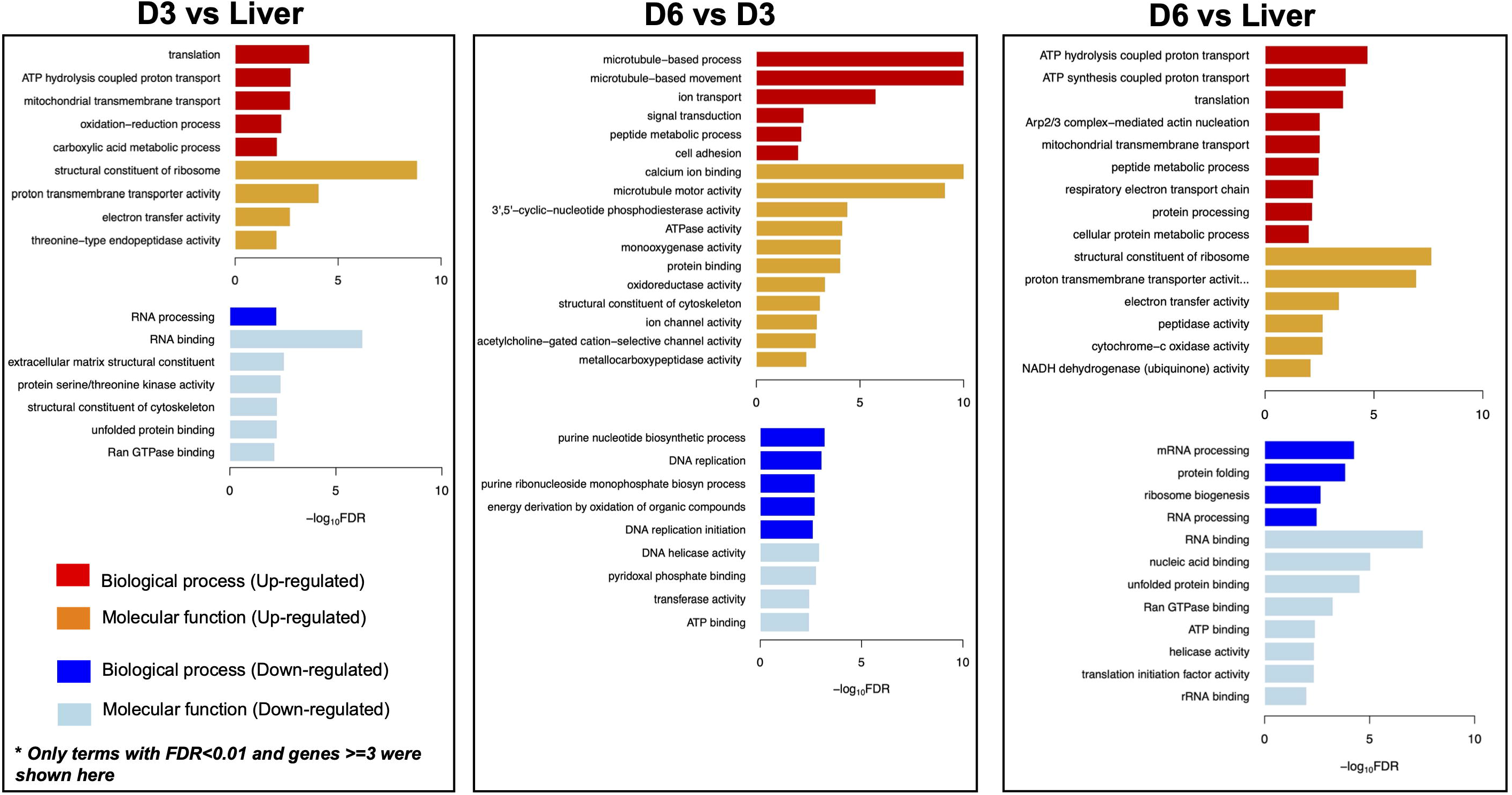
Gene Ontology (GO) enrichment in differentially expressed genes. For each comparison - D3 vs Liver; D6 vs D3; D6 vs Liver, only GO terms with FDR<0.01 and at least three genes are visualised. D3: D3 IVLE; D6: D^ IVLE; Liver: liver eggs.

Several common features between the mature egg and the miracidium were identified, and indeed the presence of fully-developed miracidia within eggshells was evident in the D6 eggs **(Figure 1B** and **Suppl. Figure S2B)**. Therefore, we asked whether a transcriptional footprint was found during the development of D6 eggs towards miracidium. To this end, we examined the top 100 genes that were previously shown to be enriched in miracidium-sporocyst (Lu and Berriman, 2018) among all developmental stages, and found that about ∼1/3 were significantly up-regulated in D6 eggs **(Suppl. Figure S7)**. The rest of the genes showed either no significant differential expression, or no expression in IVLEs **(Suppl. Table S7)**. When considering the top 200 miracidium-enriched genes, only 4.4% showed more abundance in D3 eggs, but 21.9% displayed higher expression in D6 eggs **(Suppl. Tables S4** and **S7)**, including *Sm*VAL2 (Smp_002630) and *Sm*VAL15 (Smp_070250), which were previously shown high expression in the miracidium stage (Farias et al., 2019). In addition, in D6 eggs we found an evident upregulation of *tektin* (Smp_162540) and *tubulin* (Smp_079960) genes, which are essential for microtubule assembly and physiology, key components of the ciliary machinery.

## Discussion

The schistosome egg is the main driver of chronic pathology associated with schistosomiasis (Pearce and MacDonald, 2002). In addition, it is a developmental stage that ensures the propagation of the life cycle from the definitive host to the intermediate host via the external environment. It has been recently shown that the timing of granuloma formation is actively manipulated by developing eggs, to avoid immune destruction or premature extrusion from the host (Takaki et al., 2021a). Thus, it is critical to understand the transcriptome landscape driving the egg development. Other than a handful of descriptive reports on embryogenesis (Jurberg et al., 2009) and egg secretions (Ashton et al., 2001), the number of studies focused on gene expression changes in schistosome eggs is surprisingly scarce. There is only one public transcriptome dataset for *S. mansoni* eggs and one for *S. haematobium* eggs (Young et al., 2012; Anderson et al., 2015). Mass spectrometrically-determined proteomes of the soluble egg and secreted proteins of egg, recovered from liver of *S. haematobium*-infected mice, also have been reported (Sotillo et al., 2019). In all these studies, the eggs were isolated from livers of experimentally-infected rodents and the egg transcriptomes/proteomes were compared to those of male and female adult worms aiming at identifying genes involved in host-parasite interaction. The studies did not explore changes in gene expression during embryogenesis.

Investigations of schistosome egg transcriptomes are often hindered due to intrinsic difficulties of this stage, including the presence of the eggshell and abundance of egg-derived RNases that degrade the RNA during its isolation. Here, we optimised a protocol to isolate high-quality RNA from eggs by consecutive rounds of freeze-thawing cycles. The quality of total RNA isolated from *in vitro* laid eggs (IVLE), even after 3 cycles of freeze-thawing was significantly superior to that of RNA isolated from liver eggs. Similarly, it has been reported that for biospecimens stored in tissue biobanks, such as tumour sections, the RNA quality is optimal for downstream analyses after 3 freeze-thawing cycles, but it is dramatically affected after 5 freeze-thawing cycles (Yu et al., 2017). In addition, we successfully employed the SmartSeq2 protocol (Picelli et al., 2014) to produce bulk RNA-seq data starting with few nanograms of input RNA, as has been recently shown for transcriptomic studies in *Trichuris muris* larvae (Duque-Correa et al., 2020). Repurposing single-cell RNAseq protocols to generate high-quality bulk transcriptomic data from pico- to nanograms of total RNA or from few cells (Reid et al., 2018) became a promising approach when the amount of RNA is limited.

We have previously optimised the collection of schistosome IVLE, followed and quantified their daily development (Mann et al., 2011; Yan et al., 2018). We have now produced RNA-seq data from developing eggs of *S. mansoni*. We observed drastic changes in the transcriptome profile of developing eggs and inferred functional roles of differentially expressed genes associated with embryo development (Jurberg et al., 2009). Our findings are consistent with previous descriptions of the egg development *in vitro* and the characterisation of excreted-secreted proteins by mature eggs (Ashton et al., 2001). The authors describe highly abundant protein synthesis in 3-day cultured eggs compared to freshly isolated eggs. Our findings show that the up-regulated genes in D3 vs Liver eggs and D6 vs D3 eggs are consistently associated with protein synthesis-associated processes such as translation, translational elongation, and regulation of phosphorylation. We found that genes with well-established immunomodulatory roles, e.g. *omega-1* (Fitzsimmons et al., 2005), *kappa-5* (Schramm et al., 2009) and the putative major egg antigen Sm-p40 (Stadecker and Hernandez, 1998) show a higher expression in mature eggs compared to immature eggs. However, the overall expression of these genes was much higher in eggs isolated from the host liver compared to IVLE suggesting that host factors may be required for driving their expression and/or stimulating the genesis of the subshell membrane where some of these proteins seem to be produced (Stadecker and Hernandez, 1998; Schramm et al., 2009). The eggs collected from the liver of experimentally infected rodents comprise eggs ranging from newly laid to mature eggs, several of which are dead, presumably killed by the host immune system. Thus, any comparison between the liver eggs and IVLE need to be taken cautiously. The *in vitro* culture conditions do not completely mimic the *in vivo* development of the parasite and hence, the production of fully viable eggs is limited. This is consistent with the low percentage (ranging from 10% to 15%) of IVLE that hatch fully viable and infectious miracidia (Mann et al., 2011). Recent improvements on culture conditions offer novel and informative approaches to sustain and study *in vitro* and *ex vivo* parasite development, including sexual differentiation and fecundity (Wang et al., 2019; Anisuzzaman et al., 2021; Buchter et al., 2021).

Notwithstanding the limitations of our culture system, we identified a transcriptional footprint consistent with the developmental transition from D3 to D6 egg samples and from D6 egg samples towards the miracidium. Products of several genes upregulated in D6 eggs had previously been annotated as “larval transformation proteins” during *in vitro* miracidium-to-sporocyst transformation (Wu et al., 2009), including venom allergen-like proteins (*Sm*VALs) (Farias et al., 2019). The role of *Sm*VALs is not yet confirmed, but in some parasitic nematodes, VALs are involved in mechanisms of infection establishment (Hawdon et al. 1999) and host immunomodulation (Ali et al. 2001). Whether *Sm*VALs, already upregulated in mature eggs, display similar functions during infection of the snail remains to be addressed. Proteomic approaches in *S. japonicum* identified proteins in mature eggs that are associated with miracidium motility (De Marco Verissimo et al., 2019). Similarly, we identified an upregulation of tektin (Smp_162540) and tubulin (Smp_079960) genes, both essential proteins for microtubule assembly and cilia physiology. In the D6 samples we discovered tissue-specific markers for muscle cells, nerve system, parenchymal and germ cells (Diaz Soria et al., 2020). We speculate that these genes may be involved in the specification and differentiation of diverse somatic and germinal tissues in the developing miracidia (Rawlinson et al. 2010).

In this study, we have reported the first transcriptome analysis of developing eggs from *S. mansoni*, central drivers of this major neglected tropical disease. Their transcriptomes have clear signatures of the parasite gearing up for life cycle progression, including key proteins required for the structure and motility of the miracidium and for the subsequent infection of snails. Along with highlighting proteins already known to drive egg-induced pathology, the transcriptome analysis has revealed dozens of other genes with similar profiles, not previously associated with pathogenesis but now warranting deeper investigation.

## Author contributions

G.R. conceived the project, designed the experiments, collected and cultured the *in vitro* laid eggs, interpreted the results and directed the study. Z.L. analysed the data, generated and interpreted the results. G.S. performed RNA isolation and RNA-seq. library preparation. V.O. performed the original bioinformatic analysis. KR. interpreted and discussed the results, and performed the search of *S. mansoni* orthologs of genes involved embryogenesis. P.J.B interpreted the results. M.B. provided resources and interpreted the results. Z.L. and G.R. wrote the original draft and all the authors edited the manuscript.

## Acknowledgements

We are very grateful to our colleagues at the Wellcome Sanger Institute: Simon Clare, Cordelia Brandt, Catherine McCarthy, Katherine Harcourt and Lisa Seymour for assistance and technical support with animal infections and maintenance of the *Schistosoma mansoni* life cycle; Arthur Talman and Virginia Howick for providing expertise and support with the Smart-Seq2 library preparation; Mandy Sanders for project administration. This work was funded by Wellcome (grant number 206194).

## Supplementary information

### Supplementary Figures

**Figure S1**. Total RNA isolated from different egg samples. **A**. Bioanalyzer traces of RNA preparations isolated from D3 and D6 IVLE and processed for sequencing using the Smart-Seq2 protocol. Samples and concentrations are indicated in the bottom panel. **B**. Representative bioanalyzer traces of RNA preparations isolated from IVLE and liver eggs (LE), as indicated.

**Figure S2**. Representative micrographs of D3- **(A)** and D6 **(B)** IVLE. Scale bar: 300 μm.

**Figure S3: Left**. Venn diagram indicating the number of shared/unshared upregulated genes amongst the three comparisons: D6 vs D3 IVLE (1121 genes); D3 IVLE vs liver eggs (240 genes); D6 IVLE vs liver eggs (345 genes). **Right**. Venn diagram indicating the number of shared/unshared downregulated genes amongst the three comparisons: D6 vs D3 IVLE (166 genes); D3 IVLE vs liver eggs (832 genes); D6 IVLE vs liver eggs (708 genes). The gene names and identifiers are provided in Supp. Table S3.

**Figure S4**. Pie chart indicating the differential expression of the top 200 miracidium- sporocyst enriched genes in D3 and D6 IVLE. Filtered: genes that were filtered out for differential expression analysis in DESeq2, as were detected as outlier genes; DEG: differentially expressed genes.

### Supplementary Tables

**Table S1**. RNA-Seq mapping statistics and accession numbers for all samples. Sample names refer to day of the egg collection (D3 or D6), number of biological replicate (e.g. D3_1, D3_2, D3_3), and number of technical replicates employing 3 ng (1μl - 1) or 12 ng (4μl - 4), respectively.

**Table S2**. Pearson’s correlations among the biological replicates of the *in vitro* laid eggs.

**Table S3**. Gene IDs and normalised counts for the genes numbered in the Venn diagrams in Supp. Figure S3.

**Table S4**. Lists of differentially expressed genes among liver eggs, D3, and D6 IVLEs.

**Table S5**. Lists of up-regulated genes in D6 (vs D3) IVLE that were enriched in different schistosomula cell clusters identified in (Diaz Soria et al., 2020).

**Table S6**. Enriched functions in identified differentially expressed genes (DEGs), including Gene Ontology, KEGG Pathway, and Pfam/InterPro domains.

**Table S7**. List of differentially expressed genes in D3 and D6 IVLEs enriched in miracidium-sporocyst.

### Supplementary videos

**Video S1**. Representative video showing D6 IVLE. Fully mature, motile miracidia can be seen within their eggshells. Scale bar: 50 µm.

